# Pharmacological inhibition of macrophage triglyceride biosynthesis pathways does not improve *Mycobacterium tuberculosis* control in infected mice

**DOI:** 10.1101/2024.03.06.583745

**Authors:** Jennie Ruelas Castillo, Valentina Guerrini, Darla Quijada, Styliani Karanika, Pranita Neupane, Harley Harris, Andrew Garcia, Babajide Shenkoya, Addis Yilma, Hannah Bailey, Rehan Khan, Mathangi Gopalakrishnan, Maria L. Gennaro, Petros C. Karakousis

## Abstract

Triglyceride rich macrophages (foam cells) are a hallmark of necrotic granulomas in tuberculosis, and multiple antimicrobial functions are down-regulated in these cells. In this study, we assessed the ability of two different compounds to reduce triglyceride content and intracellular burden in *Mycobacterium tuberculosis* (Mtb)-infected macrophages: A-922500 (DGATi), an inhibitor of diacylglycerol acyltransferase 1, an enzyme involved in triglyceride synthesis; and LY2584702 (p70S6Ki), an inhibitor of p70 S6 kinase, a serine/threonine kinase involved in mTORC-1dependent lipid biogenesis. Additionally, we evaluated the adjunctive activity of these inhibitors as host-directed therapies against chronic Mtb infection in C3HeB/FeJ mice. DGATi and p70S6Ki significantly reduced the lipid content and bacillary burden in Mtb-infected human monocyte-derived macrophages. In Mtb-infected mice, each inhibitor reduced the triglyceride content (P≤ 0.0001) in cells from bronchoalveolar lavage samples. Adjunctive treatment of DGATi with isoniazid and p70S6Ki monotherapy reduced the lipid droplet content (P≤ 0.05) within lung macrophages of Mtb-infected mice. However, neither inhibitor reduced the lung bacterial burden in Mtb-infected mice alone or in combination with isoniazid, and they did not alter lung inflammation. These findings provide further insights into the role of foam cells in tuberculosis pathogenesis and the utility of interventions targeting these cell populations as adjunctive host-directed therapies.

## Background

Disease caused by *Mycobacterium tuberculosis* (Mtb) is a leading cause of death from a single infectious agent worldwide, killing 1.3 million people in 2022 [1]. Host-directed therapy (HDT) offers the opportunity to enhance host defenses against Mtb infection without contributing to antibiotic resistance [2]. Foam cells represent promising targets for tuberculosis (TB) due to their integral roles in granuloma formation [3], [4], lipid metabolism [5], heightened inflammation [6], and immunomodulation [4], [7], [8]. They also play pivotal roles in the development, maintenance, and dissemination of Mtb infection [9]. Advanced granulomas exhibit a caseous necrotic core region that contributes to the establishment of a lipid-rich environment [9], [10], [11], [12]. In these granulomas, Mtb can persist and proliferate within foamy macrophages. In the necrotic core [13], the bacilli can evade phagocytosis and alter host cell death pathways [5]. Therefore, we hypothesized that modulating the lipid content within foamy macrophages may disrupt Mtb’s growth niche and compromise its survival strategies.

Triglycerides (TAG) are the most abundant neutral lipids in human TB granulomas [14], [15]. Diacylglycerol O-acyltransferase 1 (DGAT1), which converts diglycerides into triglycerides, is essential for the accumulation of TAGs in foam cells in the C3HeB/FeJ mouse model [7], which develops human-like necrotic TB lung granulomas [16]. DGAT1 also regulates the heightened inflammatory response to infection [6]. Guerrini *et al*. showed that Mtb-infected foam cells are TAG-rich and that the downstream activation of the caspases and the mammalian target of rapamycin complex 1 (mTORC1) from tumor necrosis factor receptor (TNFR) signaling leads to the accumulation of TAGs [15]. Activation of mTORC1 also promotes cellular metabolic pathways, such as glucose metabolism, protein, and TAG lipid synthesis. The 70-kDa ribosomal protein S6 kinase (p70S6K) is a major target of mTORC1 and regulates lipid biogenesis by driving the feed-forward expression of sterol regulatory element-binding transcription factor 1 (SREBP-1c) [17]. In addition, the mTORC1-p70S6K network is a known regulator of autophagy [18].

Recent studies have identified several pharmaceutical targets in the TAG biosynthesis pathway which may be repurposed as HDT agents for TB. IFN-γ-activated murine bone-marrow-derived macrophages (BMDM) infected with Mtb and treated with the DGAT1 inhibitor (T863) showed decreased lipid droplet (LD) content but unaltered Mtb replication after 24 hours [19]. However, Dawa *et al.* found that treatment with T863 led to a reduction in granuloma lipid levels and lung bacterial burden in Mtb-infected C3HeB/FeJ mice [7]. A DGAT1 inhibitor (A-922500) and rapamycin, an mTORC1 inhibitor, were shown to lower LD content in Mtb-infected human monocyte-derived macrophages (hMDMs) [15]. Finally, Mtb-infected C3HeB/FeJ mice treated with rapamycin alone or as adjunctive therapy exhibited fewer necrotic lesions and cell infiltration in the lungs than did control mice [20]. These studies underscore the potential of repurposing pharmaceutical compounds that target TAG biosynthesis to mitigate TB-associated pathology and enhance bacterial clearance.

In the present study, we evaluated the abilities of LY2584702, a highly selective inhibitor of p70S6K (hereafter referred to as p70S6Ki), rapamycin, and A-922500, a selective DGAT1 inhibitor (hereafter referred to as DGATi), to modulate TAG content and bacillary burden in Mtb-infected hMDMs. Next, we tested the effect of orally administered DGATi or p70S6Ki, as monotherapy or adjunctive therapy combined with isoniazid (INH), against chronic TB infection in C3HeB/FeJ mice. We also evaluate the potential of each of the compounds to modulate TAG content in lung macrophages and their effect on lung pathology. Overall, our findings highlight the differences in each compound’s antitubercular activities between *ex vivo* and *in vivo* models and add to the growing body of literature on the potential utility of foam cell-targeting therapies as adjunctive HDT against TB.

## Methods and Materials

### hMDMs RLU infection/treatment

hMDMs were differentiated by plastic adherence from hPBMCs, infected with the luminescent H37Rv strain at a MOI of 1 for 4 hours then washed 3 times with PBS [15]. Infected cells were incubated with fresh complete RPMI containing rapamycin, A-922500 (DGATi), LY2584702 (p70S6Ki) (all purchased from Selleckchem) or DMSO vehicle control. Three days post-infection, cells were lysed with 0.05% aqueous SDS and luminescence was measured using the GloMax 20/20 Luminometer (Promega) [15].

### hMDM lipid droplet and p62 quantification

hMDMs were infected with mCherry-expressing H37Rv infected at MOI 4 and incubated with compounds [15]. 24 hours post-infection and drug incubation, cells were washed, detached, fixed with 4% paraformaldehyde (PFA) in PBS for 45 minutes and then stained with CD11c-Bv421 for 30 minutes at 4°C. Cells were permeabilized with Cytofix/Cytoperm solution for 15 minutes at room temperature (RT) and stained with p62-APC for 30 minutes at 4°C and then with Bodipy 493/503 for 15 minutes at RT (Table S1). Images were acquired using an ImageStreamXMark II (Amnis Corporation). IDEAS software was utilized to extract cell fluorescence intensity and distribution medians for each experimental condition [15].

### *M. tuberculosis* mice infections

Wild-type Mtb H37Rv was grown in Middlebrook 7H9 broth (Difco), 10% oleic acid-albumin-dextrose-catalase (Difco), 0.2% glycerol, 0.05% Tween-80 (7H9/O/Tw at 37°C in a roller bottle to an OD_600_ of 0.8-1, followed by 1:200 dilution in 7H9/O/Tw broth. Female C3HeB/FeJ mice (6-8-week-old, Jackson Laboratory) were aerosol-infected with ∼100 bacilli of Mtb using a Glas-Col Inhalation Exposure System (Terre Haute, IN). All procedures were performed according to protocols approved by the Johns Hopkins University Institutional Animal Care and Use Committee.

### Pharmacokinetic analyses

Steady-state pharmacokinetic (PK) studies were conducted by the JHU SKCCC Analytical Pharmacology Core. Separate groups of 8-10-week-old female BALB/c mice received oral administration of one of the following: DGATi 3 mg/kg or p70S6Ki 3 mg/kg, in combination with INH 10 mg/kg, rifampin 10 mg/kg and pyrazinamide 150 mg/kg for 6 days. On the 6^th^ day, three mice were used for each time point. The PK blood samples were drawn at 0.5, 1, 2-, 4-, 8-, or 24 hours post-administration. Levels of compounds in plasma were quantified by AB Sciex, QTRAP® 5500 LC-MS/MS System instrumentation. The following PK parameters (half-life, Cmaxss, and AUClast) were determined by noncompartmental analysis using Pumas v2.5.1 software.

### Antibiotic and HDT treatment

Treatments were prepared weekly by dissolving DGATi and p70S6ki in 0.5% methylcellulose (Sigma) and INH (Sigma-Aldrich) in deionized water. Drugs were administered at the following concentrations: DGATi 30 mg/kg; p70S6Ki 30 mg/kg; INH 10 mg/kg. Mice received one of the following regimens by oral gavage 5/7 days beginning 42 days post-infection: 0.5% methylcellulose (vehicle control), INH, DGATi, p70S6Ki, INH + DGATi, or INH + p70S6Ki. Mice were euthanized and the lungs were harvested at pre-determined time points.

For colony-forming units (CFU) analysis, the superior and middle lobes of the right lung were homogenized in 2.5 mL of PBS by bead beating (2 cycles of 20s at 4500 rpm) in a Precellys® Evolution machine. Serial tenfold dilutions of lung homogenates were plated on 7H11 selective agar (BD) and incubated at 37°C for 4 weeks before counting.

### TAG quantification from bronchoalveolar lavage (BAL)

Following terminal anesthesia, BAL samples were collected by orotracheal intubation [20]. Briefly, the lungs were flushed twice with 1-ml sterile BAL buffer (PBS (Gibco), 2 nM EDTA (Corning) and 0.5% fetal bovine serum (FBS)) using a 22g cannula attached to a syringe. The lavage fluid was collected, filtered through a 70-μm cell strainer with sterile-filtered BAL complete medium (RPMI 1640, 1x glutaMAX, 1x pyruvate, 10% regular FBS), then collected by centrifugation (400 × g, 7 minutes) at 4°C. ACK lysis buffer (1 ml) was added to the pellet for 1 minute, diluted in PBS, centrifugated and resuspended in cold BAL buffer. Samples were counted with a hemocytometer and plated at 20,000 cells/well in a 96-well plate in 2-3 well replicates per animal. TAG content was quantified using the High Sensitivity Triglyceride Assay Kit (Sigma-aldrich MAK264), according to the manufacturer’s protocol, by fluorescence (λex= 535 nm/λem= 587 nm) using the FLUOstar Omega (BMG Labtech).

### Bodipy quantification for murine study

The inferior lobe of the right lung and the lateral 2/3 section of the left lung (a section was made through the hilum from the apex to the base) were placed in 2.5 mL of digestion media (RPMI 1640, Liberase 167ug/ml, DNAse I 100ug/ml). The lungs were homogenized as described above using 1 cycle of 10 seconds at 4500 rpm. The homogenate was incubated at 37°C for 15 minutes, filtered through a 70-μm cell strainer, rinsed with PBS, then centrifuged at 300g for 10 minutes at 4°C. ACK lysing buffer (1 mL) was added to the pellet for 3 minutes, then diluted with PBS, centrifuged again, and resuspended in FACS buffer (PBS + 0.5% Bovine serum albumin (Sigma-Aldrich)). Cells were stained with Zombie NIR™ for 30 minutes and washed with PBS buffer. The following anti-mouse mAbs were used for surface staining (20-minute staining): CD45-Alexa Fluor® 700, CD64-APC, Siglec-F-BV421, CD11c-PE (Table S1). Following manufacturer protocols, cells were fixed/permeabilized with buffers from Biolegend. Cells were then treated with BODIPY™ 493/503 for 30 minutes at RT in between fixing and permeabilization. Lastly, cells were washed and resuspended in FACS buffer. A BD™ LSRII flow cytometer was used. Flow data were analyzed using FlowJo Software (FlowJo 10.10.0, LLC Ashland, OR), using the gating strategies listed in Fig S3.

### Measurement of lung/body weight ratio

Lung and body weights were quantified during harvest using weight scales. For normalization purposes, the lung/body weight ratio was calculated by lung weight (g) /body weight (g) x 100.

### Quantitative analysis of lung histopathology

At the time of harvest, ∼1/3 of the left lung medial apex to base was sectioned and fixed with 4% PFA for 24 hours. Three 4-μm histological sections per lung spaced 12 μm apart were stained with hematoxylin and eosin. Digital 40X images were obtained using a brightfield microscope. Quantitative analysis was performed blinded to treatment allocation and slides pertaining to the mice. For each treatment group, a total of 21 scanned slides (3 slides per mouse) were reviewed. The lung lesion burden to whole surface area was quantified using the open-source software QuPath (https://qupath.github.io/), as described [21].

### Statistics

The differences between the treatment groups were assessed using a one-way ANOVA followed by post hoc tests (Tukey’s or Dunnett’s) or Welch’s ANOVA test (depending on the spread of the data), as stated on the figure panel description. The Prism software (GraphPad, San Diego, CA, USA) version 10.1.0 was utilized for this analysis. Results were considered statistically significant when the p-value was less than 0.05.

## Results

### Treatment of Mtb-infected human monocyte-derived macrophages with triglyceride biosynthesis inhibitors reduces lipid content and intracellular bacillary burden

To investigate the potential HDT activity of TAG biosynthesis inhibitors, we exposed hMDMs infected with a luminescent H37Rv reporter strain to a range of non-toxic doses of rapamycin, DGATi or p70S6Ki (based on EC_50_ data provided by the vendor) for optimal reduction of lipid droplet (LD) content and intrabacillary burden (data not shown). Based on these pilot studies, we selected the following compound concentrations for further study: rapamycin 0.2 nM or 0.4 nM; DGATi 30 nM; or p70S6Ki 8 nM. Three days post-infection and drug exposure, we assessed the bacillary burden in terms of relative light units (RLU) using a luminometer. LD content was quantified by staining for BODIPY493/503, a neutral lipid fluorescent dye, 24 hours post-infection and compound incubation. Relative to the vehicle, there was significant reduction of the LD content in Mtb-infected hMDMs following 24-hour exposure to rapamycin 0.2 nM (P= 0.03) or 0.4 nM (P= 0.02), but the intracellular bacillary burden was not significantly altered after 72 hours (Fig 1A). Exposure to DGATi reduced both the intracellular bacillary burden (P= 0.011) and LD (P= 0.0004) relative to vehicle (Fig 1B). Similarly, exposure of Mtb-infected hMDMs to p70S6Ki significantly reduced intracellular Mtb (P= 0.0006) and LD (P= 0.0011) relative to vehicle (Fig 1C).

**Figure 1:**
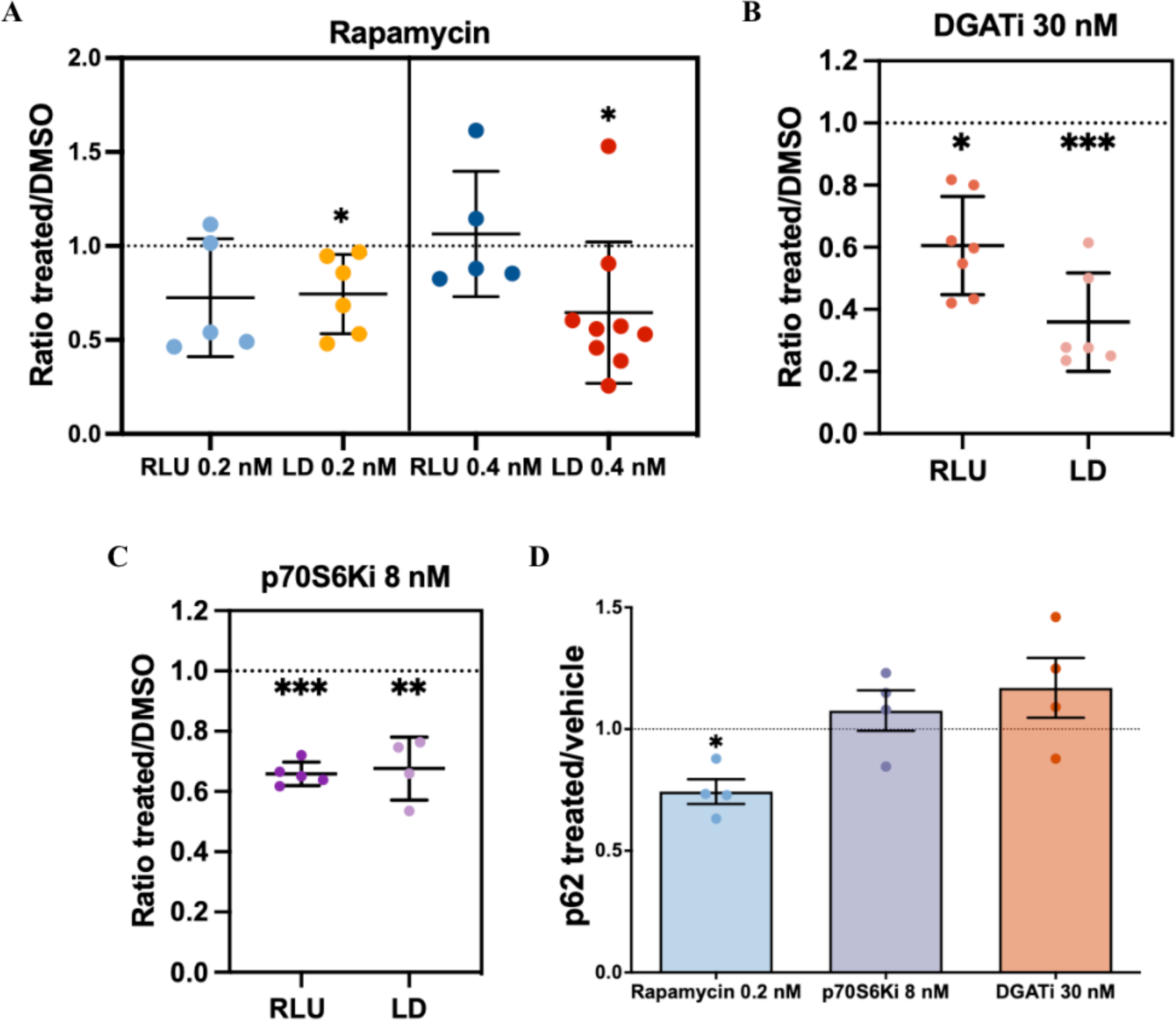
Exposure of Mtb-infected human monocyte-derived macrophages (hMDMs) to DGATi or p70S6Ki reduces lipid content and intracellular bacillary burden. hMDMs were infected with Mtb H37Rv and exposed to one of the following compounds: A) Rapamycin (0.2 or 0.4 nM); B) DGATi (30 nM); or C) p70S6Ki (8 nM). The dotted line represents values corresponding to the vehicle (DMSO) control. Lipid droplet (LD) and p62 content were quantified respectively with BODIPY493/503 and anti-p62_APC antibody staining using imaging flow cytometry, and relative luminescence units (RLU) were quantified using the GloMax 20/20 Luminometer. D) Ratio of p62 levels in Mtb-infected hMDMs treated with rapamycin, DGATi or p70S6Ki relative to vehicle-treated cells. The dotted line represents vehicle control. DGATi= diacylglycerol acyltransferase 1 inhibitor; p70S6Ki= Protein S6 Kinase inhibitor. Each dot represents individual donor results and bars show the group mean ± standard deviations. *= P < 0. 05, **= P < 0.01; ***= P < 0.001. The experiments in panels A through D consisted of four to nine donors, and each experimental condition for each donor was tested in duplicate/triplicate. A Welch’s ANOVA test performed statistical analysis for panel A and D while panels B-C were tested by one-way ANOVA followed by Dunnett’s post hoc test.

Next, since pharmacological activation of autophagy by the mTORC1-inhibitor rapamycin is known to decrease LD quantities and TAG levels [15], [22], we tested whether DGATi and p70S6Ki also induce autophagy. Using imaging flow cytometry, we quantified p62, an autophagy substrate that is used as a reporter of autophagy activity, at 24 hours post-infection and compound incubation. We found that only rapamycin 0.2 nM significantly reduced p62 (P= 0.01) relative to the vehicle (Fig 1D). Overall, these results demonstrate that DGATi and p70S6Ki are potent compounds that can reduce the intracellular Mtb burden and TAG content of infected macrophages through autophagy-independent mechanisms.

### Adjunctive therapy with DGATi or p70S6Ki reduces triglyceride levels without altering lung bacillary burden

Having established the HDT activity of p70S6Ki and DGATi in Mtb-infected hMDMs, we next tested their activity, alone and combined with INH, against chronic TB in the C3HeB/FeJ mouse model, which develops human-like, necrotic lung TB granulomas [23]. First, we established plasma pharmacokinetic profiles for DGATi 3 mg/kg and p70S6Ki 3 mg/kg with or without the adjunctive administration of the first-line regimen rifampin-isoniazid-pyrazinamide (RHZ) up to 24 hours following drug administration. We found the 24-hour steady-state drug exposures between HDT monotherapy and adjunctive therapy to be comparable (Fig S1). To maximize the potential bactericidal activity of the HDT compounds, in the primary study we administered p70S6Ki at 30 mg/kg, calculated to yield similar drug exposures to the dose previously tested in a clinical trial [24], or DGATi at 30 mg/kg, a dose shown to be well tolerated in mice [25]. We confirmed the tolerability of daily oral administration of p70S6Ki 30 mg/kg or DGATi 30 mg/kg over 14 days in C3HeB/FeJ mice (data not shown). Next, female C3HeB/FeJ mice were aerosol-infected with ∼100 CFU of Mtb-H37Rv and eight weeks later, the mice received one of the following daily (5 days/week) regimens for 3 or 6 weeks: 1) Vehicle (negative control); 2) INH 10 mg/kg (positive control); 3) DGATi 30 mg/kg; 4) DGATi 30 mg/kg + INH 10 mg/kg; 5) p70S6Ki 30 mg/kg; or 6) p70S6Ki 30 mg/kg + INH 10 mg/kg (Fig 2A). After three or six weeks of treatment, INH-treated groups had a significantly decreased lung bacillary burden compared to vehicle (P< 0.0001). Adding DGATi or p70S6Ki did not confer adjunctive activity in combination with INH (Fig 2B, Table 1). Interestingly, after six weeks of treatment, p70S6Ki alone reduced the lung bacillary burden relative to vehicle (P= 0.04) (Fig 2B, Table 1).

**Figure 2:**
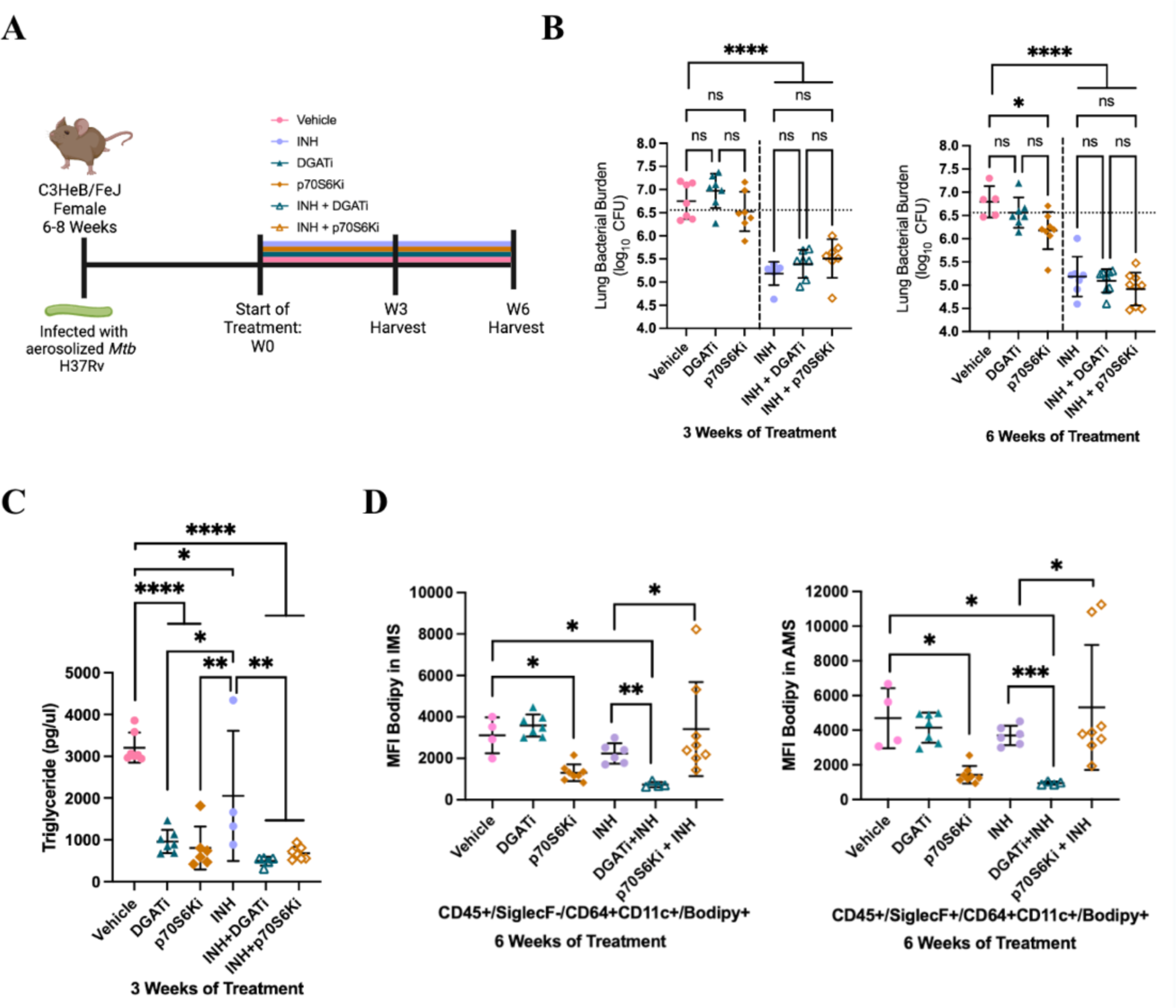
Adjunctive therapy with DGATi or p70S6Ki reduces triglyceride (TAG) levels in the lungs of Mtb-infected mice but does not alter lung bacillary burden. A) Experimental design of mouse experiments. Each line represents a different treatment group and length of treatment. C3HeB/FeJ mice were aerosol-infected with ∼100 bacilli of Mtb H37Rv. Treatment was initiated 7 weeks after aerosol infection. B) Scatterplot of lung bacillary burden after 3 or 6 weeks of treatment. The horizontal dotted line represents mean lung CFU at week 0 of treatment (W0). C) TAG concentration quantified from bronchoalveolar lavage after 3 weeks of treatment. D) Lipid droplet (LD) content determined by flow cytometry after six weeks of treatment. Individual mouse lung single-cell suspension was stained with BODIPY493/503. LD content is expressed as bodipy median fluorescence intensity (MFI). Interstitial macrophages (IMs) were represented as CD45+/SiglecF-/CD64+CD11c+/Bodipy+ and alveolar macrophages (AMs) were represented as CD45+/SiglecF-/CD64+CD11c+/Bodipy+. TAG= triglycerides; W0= week 0; W3= week 3; W6= week 6; CFU= colony-forming units; INH= isoniazid; DGATi= diacylglycerol acyltransferase 1 inhibitor; p70S6Ki= protein S6 kinase inhibitor. Each dot represents individual results and bars show the group mean ± standard deviations. *= P < 0.05; **= P < 0.001; ***= P < 0.001; ****= P < 0.0001; ns= nonsignificant. Each data point in panel B represents data from seven to nine mice per treatment group. In panel C, data represent four to 7 mice per treatment group, and in panel D there were four to eight mice per treatment group. Time course results for CFU, TAG in BAL and flow cytometry are from a single experiment. BAL TAG assay contained two to three technical replicates for each animal. Flow cytometry experiments contained two technical replicates from the same animal. Statistical analysis was performed by a one-way ANOVA followed by multiple comparison tests.

**Table 1:**
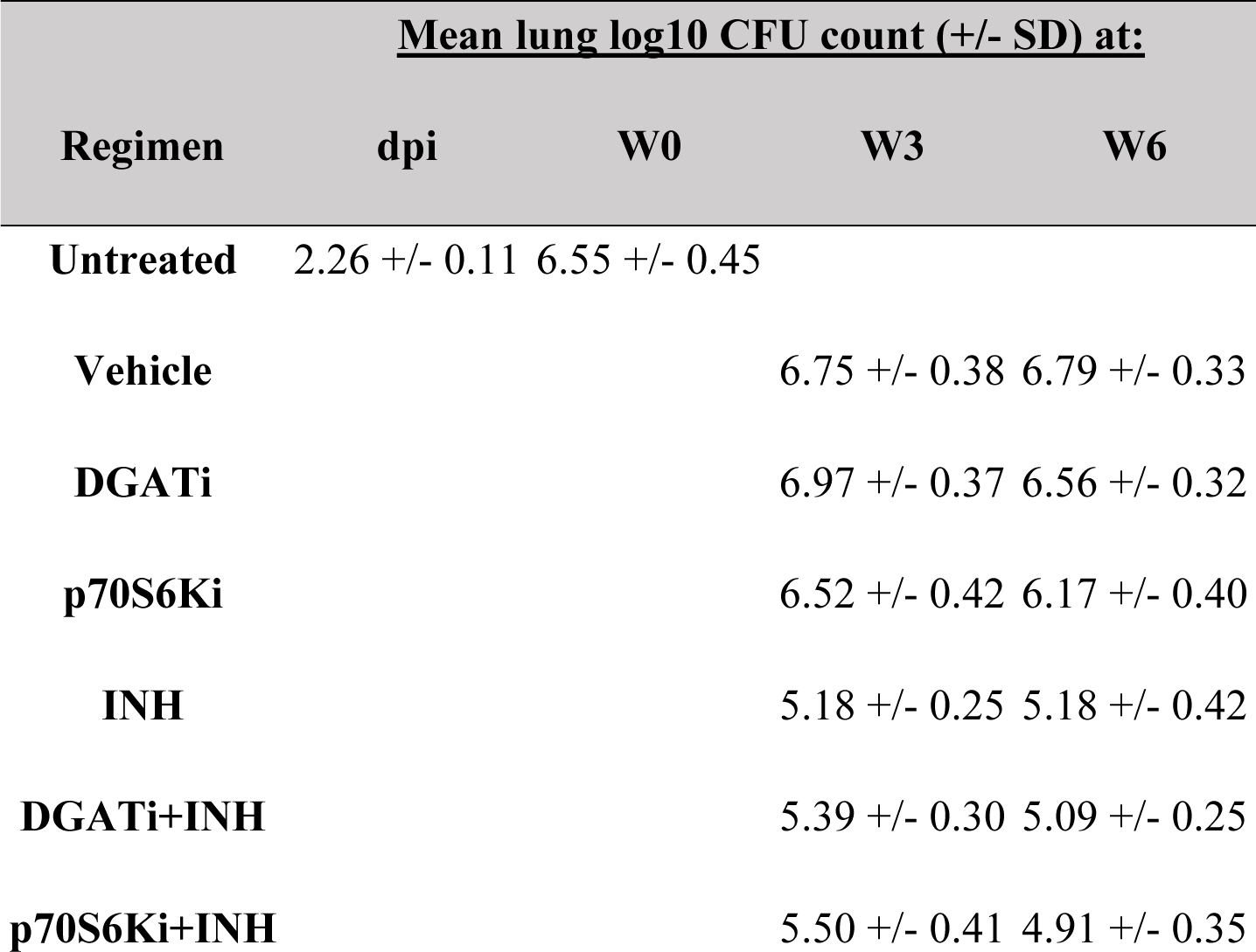
Lung Colony Forming Units (CFU) count for time course.

| Regimen | <u>Mean lung log<sub>10</sub> CFU count (+/- SD) at:</u> |  |  |  |
| --- | --- | --- | --- | --- |
|  | dpi | W0 | W3 | W6 |
| Untreated | 2.26 +/- 0.11 | 6.55 +/- 0.45 |  |  |
| Vehicle |  |  | 6.75 +/- 0.38 | 6.79 +/- 0.33 |
| DGATi |  |  | 6.97 +/- 0.37 | 6.56 +/- 0.32 |
| p70S6Ki |  |  | 6.52 +/- 0.42 | 6.17 +/- 0.40 |
| INH |  |  | 5.18 +/- 0.25 | 5.18 +/- 0.42 |
| DGATi+INH |  |  | 5.39 +/- 0.30 | 5.09 +/- 0.25 |
| p70S6Ki+INH |  |  | 5.50 +/- 0.41 | 4.91 +/- 0.35 |

In addition to the microbiological endpoints, at the time of necropsy, we harvested bronchoalveolar lavage (BAL) fluid and quantified TAG concentrations with fluorometric detection in each treatment group after 3 weeks of treatment. DGATi and p70S6Ki monotherapy and the addition of DGATi or p70S6Ki to INH significantly lowered (P< 0.0001) mean TAG concentrations in BAL samples relative to respective controls. Interestingly, INH alone also significantly lowered (P< 0.05) mean TAG concentrations (Fig 2C). To measure TAG levels in the macrophage lung cell population after six weeks of treatment, we used multiparameter flow cytometry analysis of cells stained with lipophilic dye BODIPY493/503 (bodipy). Specifically, we quantified bodipy staining in interstitial macrophages (IM) (CD45+/SiglecF-/CD64+CD11c+) and alveolar macrophages (AM) (CD45+/SiglecF+/CD64+CD11c+) (Fig S2). The percentage of live cells across all treatment groups was comparable, ranging from 85-90% mean cell viability (Fig S3A). Groups treated with vehicle, DGATi or p70S6Ki had a higher percentage of IM than AM (P= 0.0002; P< 0.0001), and (P< 0.0001, respectively). The percentage of IM and AM populations in the groups treated INH, DGATi + INH and p70S6Ki + INH did not significantly differ from each other (Fig S3B). We found that only the groups receiving p70S6Ki alone and DGATi + INH showed a significant reduction in the LD content in these cell populations relative to the vehicle (P< 0.05) (Fig 2D). Relative to INH monotherapy, adjunctive treatment with DGATi significantly decreased LD content (P< 0.01), while the addition of p70S6Ki increased LD content (P< 0.05) (Fig 2D). Taken together, these results demonstrate that these HDT agents effectively reduced TAG in the lungs of Mtb-infected mice but did not provide adjunctive antimycobacterial activity in combination with INH.

### Adjunctive treatment with triglyceride-lowering host-directed therapy agents does not reduce tuberculosis-induced lung inflammation

One of the major goals of TB HDT agents is the reduction and/or reversal of TB-induced lung damage, which can result in permanent lung dysfunction despite microbiological cure [26]. First, we analyzed the lung weight to body weight (lung/body weight) ratio to measure gross inflammation within the lungs. After three weeks of treatment, we found no significant differences between any of the treatment groups with respect to this parameter (Fig 3A). After six weeks of treatment only INH monotherapy showed a significant decrease in the lung/body ratio (Fig 3B; P= 0.006).

**Figure 3:**
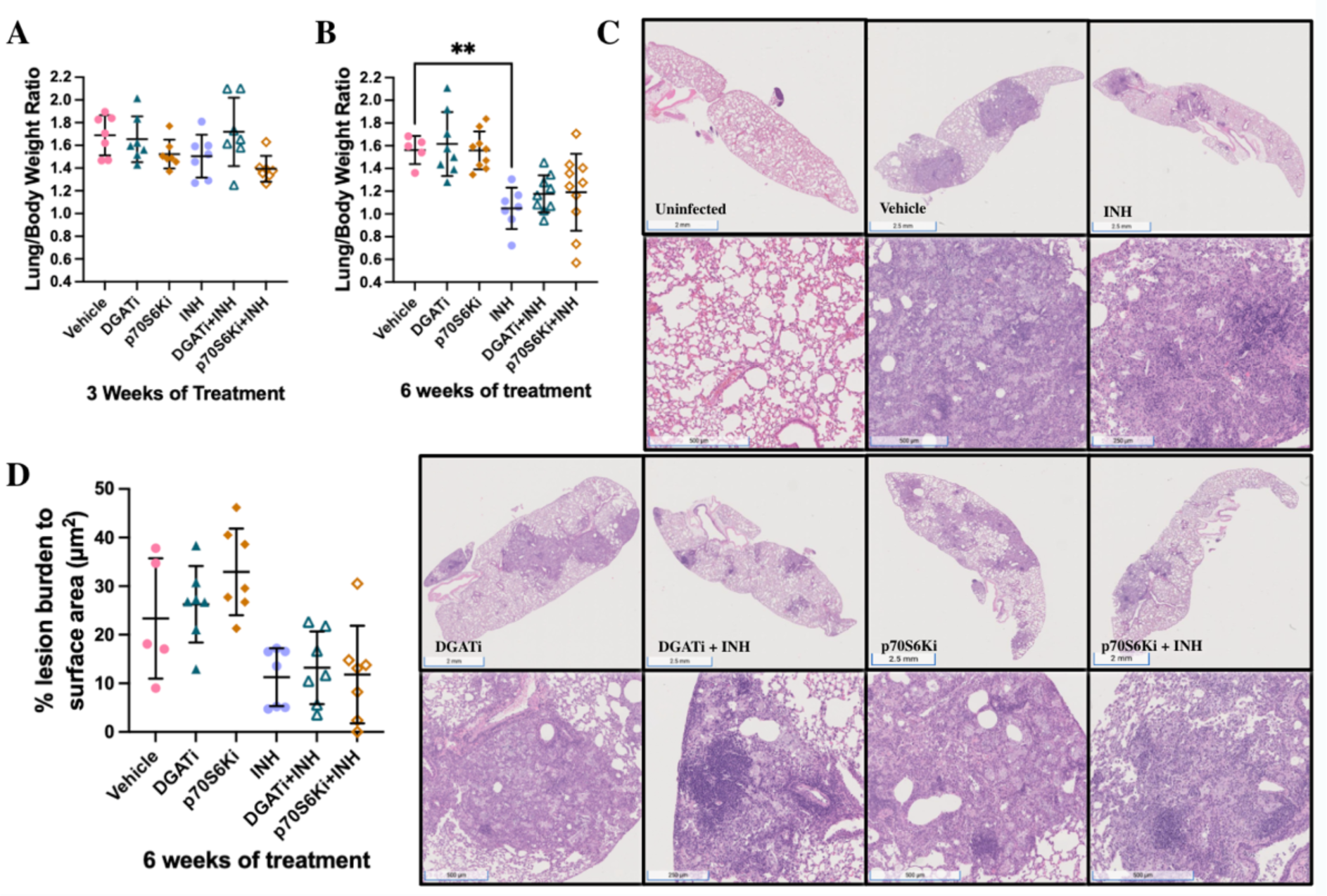
Adjunctive HDT agents do not alter TB-induced Type III lung lesions lungs or the lung/body weight ratio. Scatterplot of lung/body weight ratio after 3 (A) and 6 (B) weeks of treatment. C) Hematoxylin and eosin staining of lungs demonstrate the histopathology of different groups after 6 weeks of treatment. The top rows are representative histopathology images of the groups. The bottom rows are higher power images of the areas of inflammation within the corresponding images above. D) Quantitative analysis of histopathology after 6 weeks of treatment. INH= isoniazid; DGATi= diacylglycerol acyltransferase 1 inhibitor; p70S6Ki= protein S6 kinase inhibitor. Each group in panels A and D consisted of seven animals, panel B consisted of five to ten animals. Each dot represents individual results and bars show the group mean ± standard deviations. *= P < 0.05; **= P < 0.001; ***= P < 0.001. Statistical analysis was performed using an ordinary one-way ANOVA followed by Tukey’s multiple comparison test for panels A, B, and D.

Mtb-infected C3HeB/FeJ mice develop human-like TB lung pathology, including type I, II and III lesions, depending on the stage of disease [16]. Our histopathological analysis of lung sections from each treatment group revealed the presence only of type III lesions (Fig 3C), which are defined as inflammatory, cellular lesions composed of aggregations of lymphocytes and foamy macrophages [16]. Although there was a trend towards reduced percentage of lung surface area occupied by type III lesions in the INH-treated group, this effect did not reach statistical significance. Consistent with prior studies [7], DGAT1 inhibition was not associated with reduced TB-induced lung pathology (Fig. 3D). Similarly, mice receiving p70S6Ki alone or as adjunctive therapy in combination with INH exhibited similar lung pathology to their respective control groups (Fig. 3D).

## Discussion

HDTs offer a promising approach to complement traditional antimicrobial strategies by modulating host factors to enhance the immune response against Mtb. Targeting TAG biosynthesis could potentially disrupt the lipid-rich milieu and alter macrophage function to favor host bactericidal mechanisms. In our study, we demonstrate that in an *ex vivo* model, indirect inhibition of TAG production with DGATi (A-922500) or p70S6Ki (LY2584702), consistently reduced bacterial burden and LD within hMDMs. These inhibitors reduced the TAG content in lung macrophages of C3HeB/FeJ mice chronically infected with Mtb but failed to enhance the bactericidal activity of INH or reverse TB-induced lung inflammation. Monotherapy with p70S6Ki was the only HDT strategy found to reduce lung bacillary burden and TAG content after 6 weeks of treatment. These results suggest that adjunctive therapy with pharmacological inhibitors of macrophage TAG biosynthesis might not be an efficient TB HDT strategy in mice with chronic TB.

The current study is the first to evaluate a p70S6K inhibitor as an HDT for TB. Most HDT agents involving the mTORC1 pathway have inhibited the complex directly. Examples include rapamycin and everolimus, which were tested in Mtb models to harness their ability to induce autophagy-mediated elimination of Mtb (further reviewed in [27]). Guerrini *et al.* has previously demonstrated that rapamycin lowers LD formation within Mtb-infected macrophages [15]. In our *in vivo* study, monotherapy with the p70S6K inhibitor was found to decrease TAG content across both time points consistently, after six weeks of treatment, there was a reduction in lung bacillary burden when compared to the vehicle. In hMDMs, we found that the p70S6K inhibitor reduced LD content in an autophagy-independent manner. It is likely that the inhibition of p70S6K had a downstream effect on SREBP-1c, a key controller of *de novo* lipogenesis and TAG production in rat hepatocytes *in vivo* and *ex vivo* [28], [29], [30]. Previous studies have shown that p70S6K inhibition restricts SREBP-1c proteolytic processing [31], and decreases expression levels of lipid biosynthesis transcriptional targets [32]. In contrast, we observed no consistent reduction in the bacillary load of Mtb-infected hMDMs from different donors following exposure to rapamycin, highlighting the challenges facing HDT development across patient populations. The inhibition of DGAT1 has proven to be a validated approach for targeting macrophage TAG biosynthesis, with various pharmaceutical inhibitors demonstrating efficacy in reducing LD concentrations. Pharmaceutical inhibitors of DGAT1, such as T683 [7], [19] or A-922500 [15] reduce LD concentrations in *ex vivo* systems. Dawa *et al.* found that treatment with T863 for 15 days reduced the mean lung bacillary load in C3HeB/FeJ mice acutely infected with ∼500 CFU of Mtb Erdman [7]. However, there was a mixed response in the severity of lung infection among the treated mice. Additionally, mice treated with the inhibitor tended to have more type III lesions compared to vehicle substantial heterogeneity in lung lesion burden [7]. The discrepancies in findings between our study and those of Dawa *et al.* may be explained by key methodological differences. Although the same mouse strain was used in both studies, we infected the mice with ∼100 bacilli of the Mtb strain H37Rv and used a different DGAT1 inhibitor (A-922500). Perhaps most importantly, in our study, treatment was initiated after the establishment of chronic infection, as opposed to on day 14 after aerosol infection, as in the study by Dawa *et al*. [7]. In addition, we tested DGAT1 inhibition as an adjunctive therapeutic approach in combination with the first-line antitubercular drug INH. Most foam cells develop and localize near the central necrotic core of TB lung lesions [33], which are observed 4-6 weeks after aerosol infection [6]. Thus, it is possible that the decrease in lung bacillary burden observed with T683 therapy was due to its ability to prevent foam cell formation [7]. We believe our experimental design more closely approximates the clinical scenario, in which patients present with chronic TB disease, i.e., after the establishment of foam cells and necrotic TB granulomas, and for which patients will receive HDT not as monotherapy, but as adjunctive agents in combination with standard antitubercular drugs. Additionally, consistent with the findings of Knight *et al.*, we found no benefit from the inhibitors with respect to TB-associated pathology [19]. Our findings underscore the complex and nuanced nature of targeting TAG biosynthesis in different experimental settings. Overall, our findings suggest that pharmaceutical inhibition of macrophage TAG synthesis in macrophages might not confer a long-term host benefit to control Mtb infection.

## Author Contributions

VG, MLG, PCK conceived and designed the hMDMs studies. VG performed RLU, LD and p62 ex vivo hMDM experiments and data collection. JRC and VG performed data analysis for the hMDM studies. JRC, VG, DQ, MLG and PCK conceived and designed the time course study. SK, PN, PCK and MLG designed PK study. SK, PN conducted PK study. BS and MG assisted and conducted PK study analysis. JRC and DQ administered the therapies. JRC, DQ, SK, HP, AY, and HB performed animal harvesting, tissue sectioning and weight collection. JRC and DQ plated the homogenates of the infected lungs. JRC collected and analyzed CFU data for the time course study. JRC performed, collected data, and analyzed the TAG BAL experiment. DQ performed flow cytometry experiment. DQ and JRC collected and analyzed the flow cytometry data. SK and RK assisted with flow cytometry data visualization and analysis methodology. JRC performed data analysis for the lung/body weight ratio. AG blinded quantitative histology analysis and JRC performed the analysis. JRC, and PCK drafted the first manuscript. All authors interpreted the data and edited the manuscript.

## Acknowledgments

MLG and PCK were supported by R01HL149450 and R01AI158911 to fund these studies. We thank the Oncology Tissue Services (SKCCC) at JHU supported by the P30 CA006973 grant for their services or sectioning and imaging the H&E-stained slides. We gratefully acknowledge the assistance of Michael Urbanowski, MD, PhD, for his training and guidance in the histopathological quantification of the lung tissues. We would like to thank the New York Blood Center (Long Island City, NY, USA) for the human buffy coat samples that were obtained to make the hMDM experiments possible.

## Ethics statement

The animal study was reviewed and approved by Johns Hopkins University Institutional Animal Care and Use Committee. The animal welfare assurance number is D16-00173 (A3272-01). JHU is registered with the USDA to conduct animal research and has maintained active AAALAC accreditation since 10/4/1974.

---

**Fig S1:**
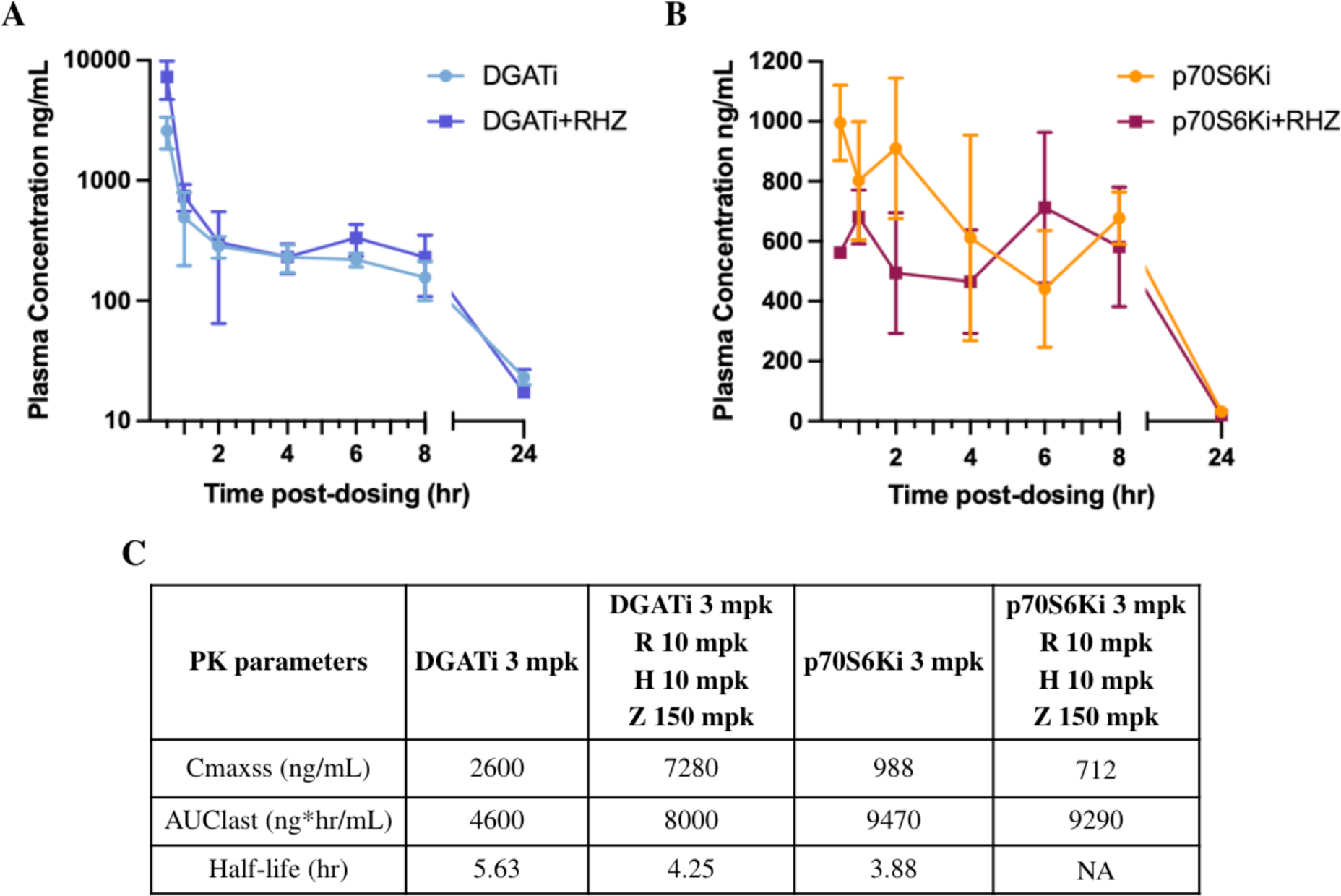
Plasma pharmacokinetic parameters for DGATi and p70S6Ki with or without rifampin-isoniazid-pyrazinamide in mice. 8-10-week-old female BALB/c mice were given DGATi, p70S6Ki, DGATi + rifampin-isoniazid-pyrazinamide (RHZ) or p70S6Ki + RHZ by oral gavage for 6 days. On the 6^th^ day, each animal contributed a single plasma pharmacokinetic (PK) sample, with at least three animals for each time point. The PK samples were drawn at 0.5, 1, 2-, 4-, 8-, or 24 hours post-administration. Plasma concentrations (ng/mL) plotted against time post-dosing up to 24 hours for animals treated with: A) DGATi or DGATi + RHZ; or B) p70S6Ki or p70S6Ki + RHZ. C) Summary of PK parameters for DGATi, p70S6Ki, DGATi +RHZ or p70S6Ki + RHZ. Cmaxss= Maximum concentration at steady state; Mpk= mg/kg; hr= hour; NA= not quantifiable. Levels of compounds in plasma were quantified by AB Sciex, QTRAP® 5500 LC-MS/MS System instrumentation. The PK parameters (half-life, Cmaxss, and AUClast) were determined by noncompartmental analysis using Pumas v2.5.1 software.

**Fig S2:**
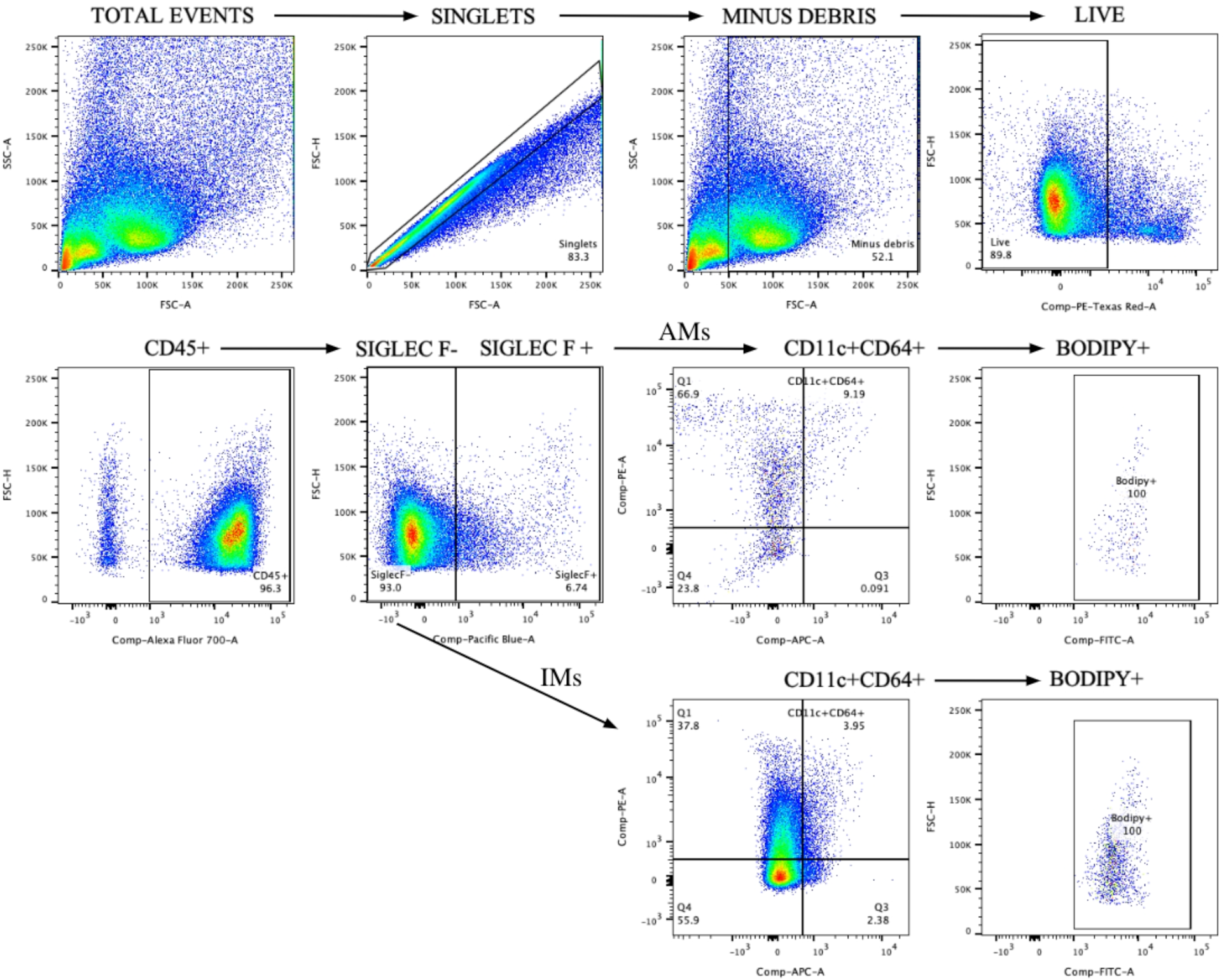
Dot plot gating strategy for identifying alveolar and interstitial macrophages by flow cytometry. Doublets and debris were excluded, and gating was performed on single live CD45+ cells. Alveolar macrophages (AMs) were classified as CD45+/ SiglecF+/CD11c+CD64+, and interstitial macrophages (IMs) were classified as CD45+/ SiglecF-/CD11c+CD64+. Neutral lipid-positive cells were identified as bodipy+ cells.

**Fig S3:**
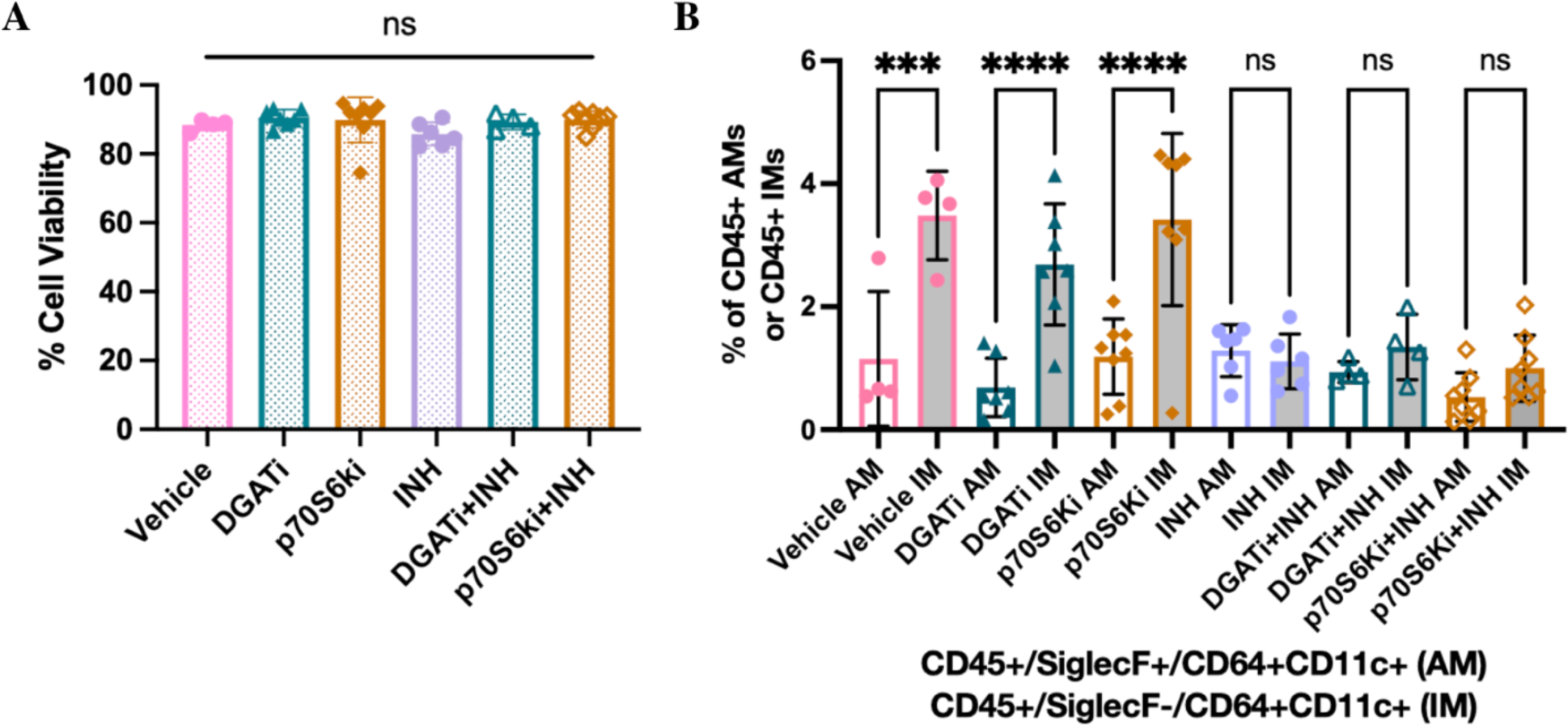
Percentage of cell viability and alveolar/interstitial macrophage populations in mouse lungs after 6 weeks of treatment. Flow cytometry results from murine lungs harvested after 6 weeks of treatment. A) Percentage of viable lung cells. Live cells were gated as singlets/minus debris/alive. B) Percentage of alveolar and interstitial macrophages populations gated from CD45+ cells. Alveolar macrophages (AMs) were classified as CD45+/SiglecF+/CD11c+CD64+, and interstitial macrophages (IMs) were classified as CD45+/SiglecF-/CD11c+CD64+. INH= isoniazid; DGATi= diacylglycerol acyltransferase 1 inhibitor; p70S6Ki= protein S6 kinase inhibitor. For panels A and B, each dot represents individual results and bars show the group mean ± standard deviations. Each group consisted of four to eight mice. Ns= nonsignificant; ***= P < 0.001; ****= P < 0.0001. Statistical analysis was performed by an ordinary one-way ANOVA followed by Tukey’s multiple comparison test for panels A and B.

**Table S1:** hMDM and mouse flow cytometry study reagents and antibodies.

| Reagents/Antibodies | Source |
| --- | --- |
| CD11c-Bv421 | BD |
| Cytofix/Cytoperm solution | BD |
| p62-APC | ABCAM |
| BODIPY™ 493/503 | Invitrogen Cat. No.: D3922 |
| Zombie NIR™ Fixable Viability Kit | Biolegend Cat. No.: 423105 |
| Alexa Fluor® 700 anti-mouse CD45 Antibody | Biolegend Cat. No.: 103127 |
| APC anti-mouse CD64 (FcγRI) Antibody | Biolegend Cat. No.: 139305 |
| Siglec-F Rat anti-Mouse, BV421, Clone: E50-2440 | BD Horizon Cat. No.: BDB565934 |
| PE anti-mouse CD11c Antibody | Biolegend Cat. No.: 117307 |
| Biolegend intracellular fixation/permeabilization set | Biolegend Cat. No.: 421002 |

